# Paranemic cohesion of DNA molecules in different counter ions at room and physiological temperatures

**DOI:** 10.1101/2025.08.21.671517

**Authors:** Lauren Anderson, Hannah Talbot, Arun Richard Chandrasekaran

**Affiliations:** Department of Nanoscale Science and Engineering, University at Albany, State University of New York, Albany, NY, USA; Department of Biological Sciences, University at Albany, State University of New York, Albany, NY, USA; The RNA Institute, University at Albany, State University of New York, Albany, NY, USA

## Abstract

Isothermal assembly allows the construction of DNA nanostructures at constant moderate temperatures. Further, assembly of DNA nanostructures in different counter ions expands their utility in various applications. In this work, we demonstrate the isothermal assembly of paranemic crossover (PX) DNA motifs in magnesium (Mg^2+^), calcium (Ca^2+^) and strontium (Sr^2+^) at 20 °C and 37 °C. Further, we study how differences in design affect the assembly of 4-stranded and 2-stranded PX molecules.

---

Paranemic crossover (PX) DNA is a multi-stranded, multi-crossover motif used in the assembly of DNA objects,^1^ devices,^2,3^ arrays^4,5^ and origami.^6,7^ PX DNA consists of two adjacent and connected double helical DNA domains, formed by creating crossovers between strands of the same polarity at every possible point between the two helices.^6,7^ Each duplex domain contains alternating wide or narrow groove separation flanking the central dyad axis of the structure, with the helical repeat of a PX molecule (∼22 bp) being twice as that of conventional B-DNA. PX DNA can be formed from 4 different strands or 2 strands that are looped on one or both ends (**Figure 1**). PX DNA is typically assembled in magnesium-containing buffers using a thermal annealing protocol. In the case of paranemic cohesion of DNA objects, two half-PX molecules have been shown to cohere at 37 °C at high magnesium concentrations to yield a full PX molecule.^8^ Recent progress in DNA nanotechnology, including our own recent work, has shown the isothermal assembly of several DNA nanostructures by incubation at constant temperatures instead of a thermal annealing protocol.^9–15^ Further, we recently demonstrated that DNA nanostructures can be assembled in different counter ions instead of the typically used magnesium.^9,16,17^ In this work, we expand the isothermal assembly method to PX DNA, showing isothermal assembly in magnesium (Mg^2+^), calcium (Ca^2+^) and strontium (Sr^2+^) for 4-stranded and 2-stranded PX molecules.

**Figure 1.**
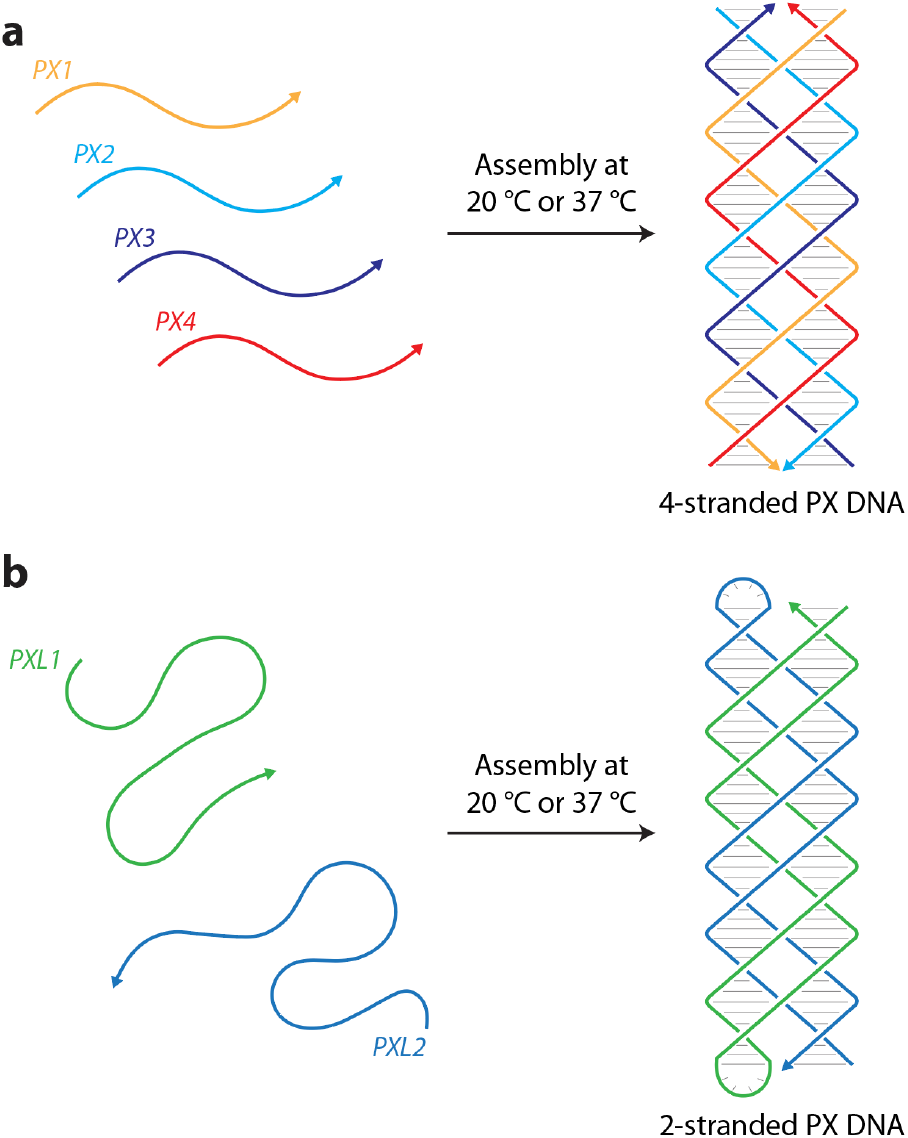
Design of PX molecules. (a) A 4-stranded PX molecule assembled from four 38-nt long oligonucleotides. (b) A 2-stranded PX molecule assembled from two 80-nt long oligonucleotides containing a 4-thymine loop in the middle.

The PX structure we chose contains 6 nucleotides in the major groove and 5 nucleotides in the minor groove, known to be the most stable among PX complexes.^6,7,18^ The 4-stranded version contains four 38-nucleotide strands that hybridize together to form the PX molecule (**Figure 1a** and **Table 1**). For the 2-stranded PX, we chose the same set of sequences but added a loop containing four thymines to connect two strands on either side of the molecule, resulting in two 80-nucleotide component strands (**Figure 1b** and **Table 1**).

**Table 1.**
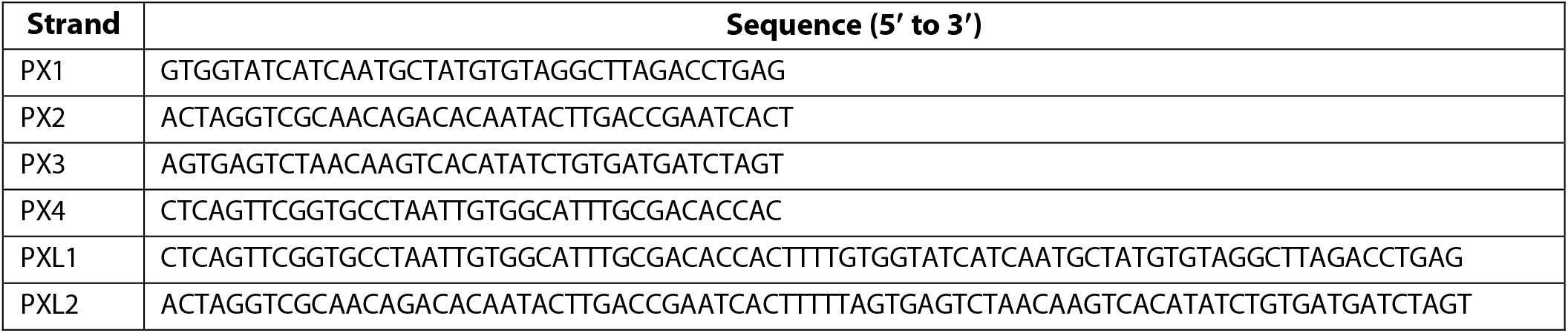
Sequences used in this study.

We first tested the effect of different Mg^2+^ concentrations on isothermal PX DNA assembly. For potential practical applications, we chose room temperature (20 °C) and physiological temperature (37 °C) as the assembly temperatures. We mixed the component strands in tris-acetate-EDTA (TAE) buffer containing 12.5 to 125 mM Mg^2+^ and incubated the samples at constant temperatures of 20 °C or 37 °C for 1 hour. We analyzed the incubated samples using non-denaturing polyacrylamide gel electrophoresis and observed assembly of the structure in both temperatures, and at all tested ion concentrations. To evaluate differences in assembly yields, we quantified the band intensity corresponding to the structure, and defined the yield as the fraction of the target band intensity among all the products in the gel lane (**Figure 2a**). To compare isothermal assembly, we normalized this yield to the assembly yield of the structure annealed in TAE buffer containing 12.5 mM Mg^2+^, the an often used buffer for such DNA motifs. We observed an increase in the assembly yield with increasing Mg^2+^ concentration, with assembly yields at 37 °C being slightly higher than those at 20 °C. When isothermally assembled, the highest assembly yields in 125 mM Mg^2+^ were ∼89% and ∼104% for 20 °C and 37 °C respectively when compared to the annealed control. We then expanded the study to Ca^2+^ and Sr^2+^ ions by using the same TAE buffer but substituting Mg^2+^ for 12.5 to 125 mM Ca^2+^ or Sr^2+^. Isothermal assembly in Ca^2+^ was similar to that of Mg^2+^, with ∼87% and ∼104% yields in 125 mM Ca^2+^ at 20 °C and 37 °C respectively. Assembly yield in Sr^2+^ ion was slightly lower compared to Mg^2+^ and Ca^2+^, with ∼81% at 20 °C and ∼89% at 37 °C in 125 mM ion concentration.

**Figure 2.**
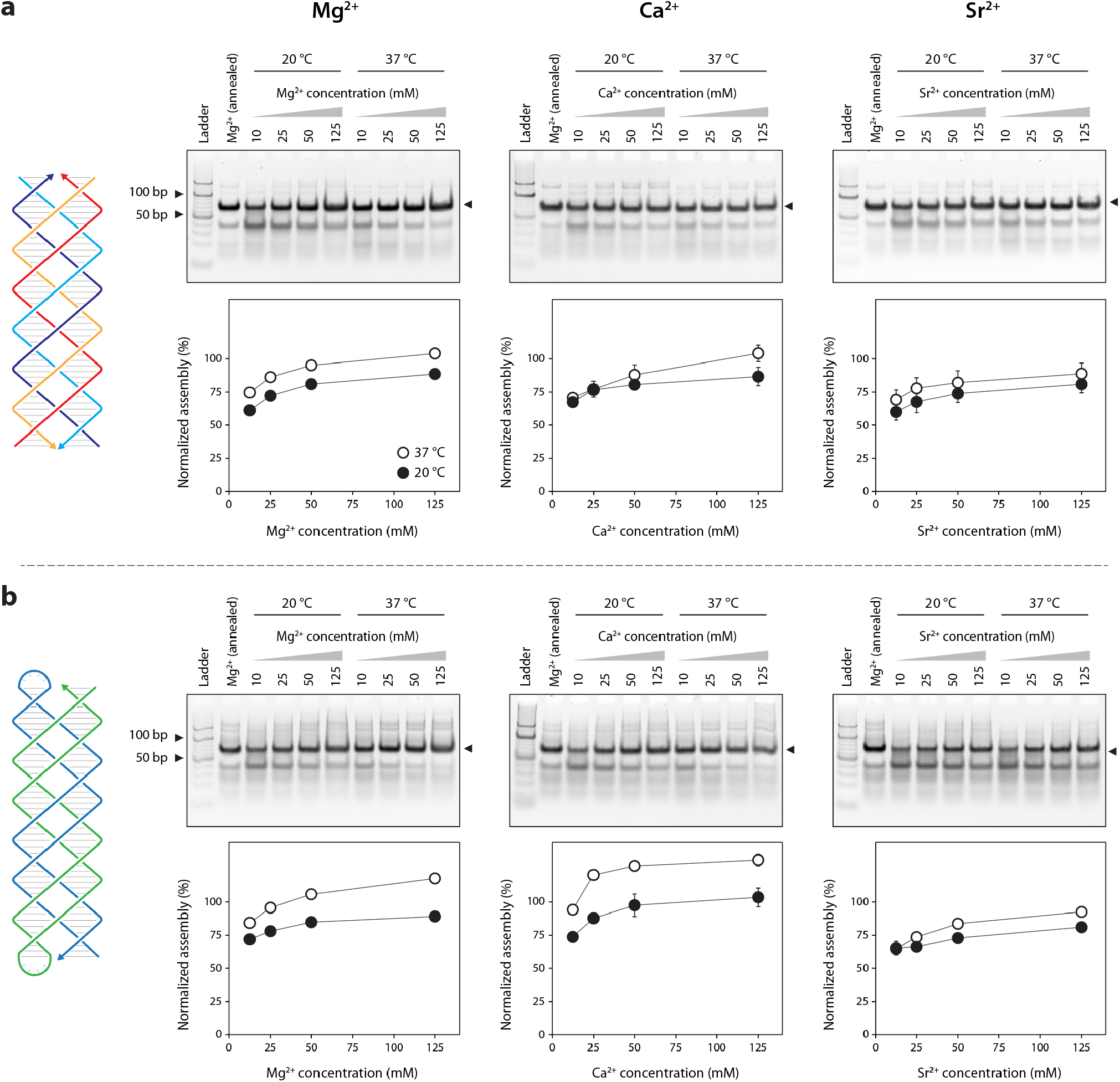
Isothermal assembly of PX molecules. (a) Assembly of the 4-stranded PX molecule in TAE buffer containing different concentrations of Mg^2+^, Ca^2+^ and Sr^2+^ at a constant temperature of 20 °C or 37 °C. (b) Assembly of the 2-stranded PX molecule in TAE buffer containing different concentrations of Mg^2+^, Ca^2+^ and Sr^2+^ at a constant temperature of 20 °C or 37 °C. Assembly yields are normalized to the yield of the specific structures assembled in TAE buffer containing 12.5 mM Mg^2+^ using a thermal annealing protocol.

We next performed a similar set of experiments to analyze the isothermal assembly of the 2-stranded PX DNA (**Figure 2b**). While the trend of assembly yields was similar for both the 4-stranded and the 2-stranded PX molecules with regards to ion concentration, we observed slight differences between the two structures. In Mg^2+^, isothermal assembly at 37 °C in 25 mM Mg^2+^ was comparable to the annealed structure (∼96% normalized yield), whereas the 4-stranded PX had a similar yield only with 50 mM Mg^2+^. Similarly, assembly yield of the 2-stranded PX in 12.5 mM Ca^2+^ at 37 °C (∼94%) was comparable to the annealed structure while surpassing the yield of the annealed sample in Mg^2+^ when isothermally assembled in 25, 50 and 125 mM Ca^2+^ (120-132% normalized yields). These yields were higher than the assembly yields of the 4-stranded PX DNA in Ca^2+^. In contrast, the assembly yields in Sr^2+^ was similar for the 4-stranded and the 2-stranded structures at both isothermal temperatures. In the 4-stranded PX, all four strands hybridize to form the structure whereas the strands in the 2-stranded PX structure require intramolecular association to form a half-PX followed by the formation of the full PX molecule. These results indicate that differences in inter- and intramolecular association of strands play a role in the assembly of DNA motifs at specific temperatures.

Overall, we show that isothermal paranemic cohesion of DNA molecules is possible in different counter ions with a short incubation time. Further, achieving this assembly at room temperature and physiological temperature makes this work adaptable for DNA nanotechnology applications in ambient environments. PX cohesion has been shown to be stronger than sticky-ended cohesion,^6^ and the ability to connect nanostructures at room temperature allows the interfacing of DNA nanostructures and arrays with polymer and silica based structures for materials science applications. Assembly under physiological conditions shown here, in combination with the exceptional biostability of PX molecules,^19,20^ could be useful in biological applications. Further, reconfiguration of DNA nanostructures held by PX cohesion can be achieved at environmental temperatures using this strategy. Combined with prior work on strand displacement based PX devices^2,3^ and our earlier work on mismatch and enzyme based operation of PX devices,^21,22^ isothermal assembly allows both the construction and operation of such devices at one constant temperature.

## Methods

### Preparation of DNA complexes

DNA strands were purchased from Integrated DNA Technologies (IDT). For different metal ions, the chemicals used were magnesium acetate tetrahydrate, calcium chloride dihydrate and strontium chloride (Sigma-Aldrich). For control DNA complexes annealed in Mg^2+^, component DNA strands were mixed in 1× TAE buffer containing 40 mM tris base (pH 8.0), 20 mM acetic acid, 2 mM EDTA, and 12.5 mM magnesium acetate. For samples annealed in other ions, component DNA strands were mixed in 1×?TAE buffer containing the specified amounts of the corresponding metal salt. Control samples were annealed in a thermal cycler with the following steps: 90 °C for 3 minutes, 65 °C for 20 minutes, 45 °C for 20 minutes, 37 °C for 30 minutes, 20 °C for 30 minutes, and the solution was then cooled to 4 °C. For isothermal assembly, component DNA strands were mixed in 1× TAE containing the specified amounts of metal salt and incubated at 20 °C or 37 °C for 1 hour.

### Non-denaturing polyacrylamide gel electrophoresis

DNA samples were mixed with 10× loading dye containing bromophenol blue and glycerol and loaded in polyacrylamide gels (19:1 acrylamide solution, National Diagnostics) prepared in 1× TAE buffer containing 12.5 mM Mg^2+^. Gels were run at 4 °C with 1× TAE-Mg^2+^ as the running buffer and stained with 0.5× GelRed (Biotium) in water for 20 min in dark and destained in water for 10 min. Gels were imaged on a Bio-Rad Gel Doc XR+ imager using the default settings for GelRed with UV illumination and analyzed using ImageLab software (Bio-Rad). Images were typically taken at multiple exposures to facilitate accurate quantification. For each gel, quantification was done using the highest-exposure image that did not contain saturated pixels in the bands. The assembly yield was quantified as the fraction of the intensity corresponding to the band of interest compared to the total intensity of all the bands in the lane. This yield was then normalized to the assembly yield of the structure prepared by annealing in 1× TAE buffer containing 12.5 mM Mg^2+^.

## Author contributions

L.A. performed experiments and analyzed data. H.T. designed experiments, performed experiments, analyzed data and edited the paper. A.R.C. conceptualized and supervised the project, designed experiments, analyzed data, visualized data and wrote the paper.

## Acknowledgments

Research reported in this publication was supported by the National Institutes of Health (NIH) through National Institute of General Medical Sciences (NIGMS) under award number R35GM150672 to A.R.C. This manuscript is the result of funding in whole or in part by the National Institutes of Health (NIH). It is subject to the NIH Public Access Policy. Through acceptance of this federal funding, NIH has been given a right to make this manuscript publicly available in PubMed Central upon the Official Date of Publication, as defined by NIH.

